# Genome-Wide Identification and Expression Pattern of the *N*-acetylserotonin deacetylase (ASDAC) Gene Family in Orchidaceae

**DOI:** 10.1101/2024.05.09.592847

**Authors:** Enda Sun, Erqiang Zhao, Qianqian Li, wenxiu lu, JiaQI Jin, Yingjia Li, Chen Yang, Tingying Chen, Zongmin Mou, Dake Zhao

## Abstract

Orchids are a kind of horticultural plant with highly ornamental and medical value. *N*-acetylserotonin deacetylase (ASDAC) is the only reverse enzyme of the melatonin biosynthesis pathway, and plays an important role in regulating the balance of melatonin. Melatonin as a multifunctional molecule, is typically involved in plant growth and development regulation, as well as abiotic stress tolerance. Here, we aimed at identifying *ASDAC* genes from the orchid genome to provide valuable information for further study of the role of melatonin in orchids.

In this study, a total of 7 *ASDAC* genes were identified from the 7 orchid genome with one member in each species. The 7 orchid *ASDACs* have an HDAC functional domain, and cluster together with functionally confirmed *OsHDAC10* and *AtHDAC14*, it shows that these members may have function of *N*-acetylserotonin deacetylase. Furthermore, based on the phylogenetic, motif, and gene structure analysis, the same cluster’s orchid *ASDAC* or *ASDAC*-like genes generally contained similar introns and motifs, suggesting the distribution pattern of exons/introns and motifs were strongly related to phylogeny on an evolutionary basis. Interestingly, homologous genes of *OsHDAC10* and *AtHDAC14* in *Gastrodia elata* have low homology and not cluster together with rice and *A.thaliana ASDACs*, showing that *ASDAC* gene family may lost in the holomycoheterotrophic orchids. The Ka/Ks ratios of *ASDAC* gene pairs from lower plant to higher plant were less than one, suggested that *ASDAC* genes have undergone purifying selection during the evolution process. *Cis*-acting element analysis results showed that the promoter regions of orchid *ASDAC* genes contained plant growth and development, phytohormone, and stress-responsive elements. Moreover, most orchid *ASDACs* were expressed in the roots, stems, leaves, flowers, and seeds. Combined *cis*-acting element and tissue expression analysis, indicating orchid *ASDAC* genes are involved in melatonin regulation of growth and development, as well as melatonin responding to various stresses in orchids. These findings of orchid *ASDAC* genes may provide valuable information for further study of the role of melatonin in orchids.

## 1. INTRODUCTION

Orchidaceae is one of the largest families of angiosperms with approximately 28,000 accepted species^[1, 2]^. As an important horticultural plant, the family has highly economic value and is widely used in the medical, ornamental, and cosmetics industries^[3]^. Moreover, the orchids possess an extraordinary lifestyle diversity and grow in almost all terrestrial habitats^[4, 5]^. To live in these habitats, orchids usually encounter abiotic stress (drought, salt, and light)^[3]^. Melatonin (*N*-acetyl-5-methoxytryptamine), a small indole compound, plays important roles in plant growth and development such as seed germination^[6, 7]^, root development^[8]^, leaf senescence^[9]^, stem strength^[10]^, and flower development^[11]^. Additionally, melatonin is involved in multiple abiotic stress tolerance, containing drought, waterlogging, salt, cold, heat, and UV-B^[12, 13]^. However, melatonin mediated growth regulation and stress response of orchids are still unknown.

*N*-acetylserotonin deacetylase (ASDAC), the only reverse enzyme of the melatonin biosynthesis pathway, plays an imperative role in regulating the balance of melatonin^[14]^. In this way, melatonin may be maintained at an optimal level, which is essential for plant growth and development, as well as response to abiotic stress. Hence, to understand the function of melatonin, we conducted a genome-wide analysis of *ASDAC* genes in Orchidaceae and made comprehensive assessments of the chromosomal localization, phylogenetic relationship, motif distribution, gene structure, Ka/Ks ratios, *cis*-active regulatory elements, and tissue-specific expression. These results provide insight into the foundation for further study of the function of melatonin in Orchidaceae.

## 2. MATERIALS AND METHODS

### 2.1 Identification and Chromosomal Localization of ASDAC Family Genes in Orchidaceae

The previously identified rice (OsHDAC10: Accession no. AK072557) and *A. thaliana* (AtHDAC14: Accession no. At4g33470) ASDAC protein sequences were used as a query to search the *Apostasia shenzhenica, Phalaenopsis equestris, Dendrobium catenatum, Phalaenopsis aphrodite, Cymbidium ensifolium, Gastrodia elata, Dendrobium chrysotoxum, Dendrobium nobile*, and *Vanilla planifolia* genomic databases by the National Center for Biotechnology Information (NCBI, https://www.ncbi.nlm.nih.gov/), OrchidBase 2.0, and Orchidstra 2.0 online websites^[14-16]^. Candidate sequences were selected with an E value less than 10^−30^, and submitted to the Simple Modular Architecture Research Tool (SMART, http://smart.embl-heidelberg.de/) database^[17]^ to validate the ASDAC domains. After removing protein sequences without a typical ASDAC conserved domain, the candidate orchid ASDAC proteins were finally confirmed. The molecular weight (Mw) and isoelectric points (pI) of the candidate ASDAC proteins were predicted by the ExPASy database (https://web.expasy.org/compute_pi/)^[18]^, and the subcellular localization was predicted using the WoLF PSORT tool (https://www.genscript.com/wolf-psort.html?src=leftbar)^[19]^.

The chromosome location of ASDAC proteins were retrieved from the GFF3 file of *D. chrysotoxum* and *D. nobile* genome^[20, 21]^, and then mapped the DchHDACs and DnHDACs onto the specific chromosomes using the Mapgene2chrom v2.1 online tool (http://mg2c.iask.in/mg2c_v2.1/)^[22]^.

### 2.2 Multiple Sequence Alignment and Phylogenetic Analysis

The full-length amino acid sequences of candidate ASDAC proteins were aligned with the protein sequences of *OsHDAC10* and *AtHDAC14* using DNAMAN 8 tool. In order to study the evolutionary relationships of orchid *ASDACs*, the sequences of other putative *ASDACs* were retrieved from the NCBI and Phytozome (https://phytozome-next.jgi.doe.gov/) database^[23]^ (**Supplementary Table S3 and S4**). The phylogenetic tree was constructed using the maximum likelihood (ML) method with default parameters by PhyloSuite software^[24]^. Next, the phylogenetic tree was visualized using the Fig Tree and iTOL (https://itol.embl.de/) tool^[25]^.

### 2.3 Gene Structure and Conserved Motif Analysis

The exon-intron structures of orchid *ASDACs* and *ASDAC* genes from various taxa (containing archaea, algae, mosses, ferns, and spermatophyte) were identified using Gene Structure Display Server 2.0 (GSDS 2.0, http://gsds.gao-lab.org/) online tools^[26]^. The conserved motifs of ASDAC proteins were analyzed by Multiple Em for Motif Elicitation (MEME, https://meme-suite.org/meme/) software^[27]^ with the following parameters: maximum number of motifs-10, and the other parameters were default. The results from MEME were visualized using TBtools^[28]^.

### 2.4 Calculation of Ka, Ks, and Ka/Ks

DNAMAN 8 software was used to calculate the identity of *ASDAC* gene pairs. Online tool PAL2NAL (http://www.bork.embl.de/pal2nal/)^[29]^ was used to calculate Ka (non-synonymous substitution rate), Ks (Synonymous substitution rate), and Ka/Ks ratios. The default parameters were employed in the above-mentioned analysis.

### 2.5 Cis-Acting Regulatory Elements Analysis in the Promoters

The 2000 bp sequence upstream of the start codon of *A. shenzhenica, P. equestris, D. catenatum, P. aphrodite*, and *D. nobile ASDAC* genes were extracted using the TBtools^[28]^. The potential *cis*-elements of orchid *ASDACs* were searched in the PlantCARE database (http://bioinformatics.psb.ugent.be/webtools/plantcare/html/)^[30]^. Finally, data arrangement and visualized using the Excel and TBtools^[28]^.

### 2.6 Gene Expression Analysis

To analyze the expression patterns of *ASDAC* genes in Orchidaceae, the expression data in various plant tissues of *A. shenzhenica* (inflorescence, leaf, root, seed, stem, tuber, and pollinium), *P. equestris* (sepal, petal, labellum, pollinium, gynostemium, floral stalk, leaf, root, and seed), *D. catenatum* (flower bud, sepal, labellum, pollinium, gynostemium, stem, leaf, root, green root tip, and white part root), *P. aphrodite* (fully open flower, large buds, small buds, leaf, root, long stalk (1.5∼3 cm), short stalk (≤ 1 cm), pollinium, and seed), *C. ensifolium* (pooled 4 stage of bud and mature flower), and *V. planifolia* (leaf, stem, and root) were downloaded from OrchidBase 2.0 and Orchidstra 2.0 database^[15, 16]^.

## 3. RESULTS

### 3.1 Primary Identification of Orchid *ASDAC* Genes

To identify the presumed orchid *ASDAC* genes, rice and *A. thaliana* ASDAC proteins were used as a query to search for ASDAC proteins in orchid protein files. After confirming in the SMART databases, a total of 34 putative *ASDAC* genes were obtained. Among them, 4 members of ASDAC family were identified in the genome of *A. shenzhenica* (named as *AsHDAC1* to *AsHDAC4*), *P. equestris* (named as *PeHDAC1* to *PeHDAC4*), *D. catenatum* (named as *DcaHDAC1* to *DcaHDAC4*), *P. aphrodite* (named as *PaHDAC1* to *PaHDAC4*), *D. chrysotoxum* (named as *DchHDAC1* and *DchHDAC4*), and *V. planifolia* (named as *VpHDAC1* and *VpHDAC4*), respectively. 5 members of ASDAC family were identified in the genome of *C. ensifolium* (named as *CeHDAC1* to *CeHDAC5*), 3 members of ASDAC family were identified in the genome of *D. nobile* (named as *DnHDAC1* to *DnHDAC3*), and 2 members of ASDAC family were identified in the genome of *G. elata* (named as *GeHDAC1* and *GeHDAC2*). Sequences including coding sequence (CDS) and protein sequences of candidate orchid *ASDAC* genes are listed in **Supplementary Table S1**. The basic features of the candidate *ASDAC* genes were predicted, details concerning the gene name, gene locus, open reading frame (ORF) length, protein length, molecular weights (Mw), isoelectric points (pI), and protein subcellular localization (**Table 1)**. The length of the candidate ASDAC proteins ranged from 292 to 694 amino acids, the ORF lengths ranged from 879 to 2085 bp, the Mw ranged from 31496.4 to 77383.33 Da, and the pI ranged from 5.03 to 8.69. Moreover, the prediction of protein subcellular localization revealed that 22 orchid ASDAC proteins resided in the cytoplasm region, 6 ASDAC proteins resided in the chloroplast region, 5 ASDAC proteins resided in the nucleus region, and only one ASDAC protein resided in the mitochondrion region.

**TABLE 1.** A list of *ASDAC* genes in *A. shenzhenica, P. equestris, D. catenatum, P. aphrodite C. ensifolium, G. elata, D. chrysotoxum, D. nobile*, and *V. planifolia*, and their characteristics.

| Name | Gene locus | ORF length<br>(bp) | Length<br>(aa) | Mw (Da) | pI | Subcellular<br>localization |
| --- | --- | --- | --- | --- | --- | --- |
| AsHDAC1 | Ash000351 | 1290 | 429 | 46852.51 | 6.16 | mitochondrion |
| AsHDAC2 | Ash010680 | 1146 | 381 | 41923.33 | 5.75 | nucleus |
| AsHDAC3 | Ash015221 | 2061 | 686 | 76413.68 | 5.29 | cytoplasm |
| AsHDAC4 | Ash002246 | 1863 | 620 | 68519.43 | 5.86 | nucleus |
| PeHDAC1 | Peq001683 | 1458 | 485 | 52725.84 | 6.38 | cytoplasm |
| PeHDAC2 | Peq002924 | 2031 | 676 | 75180.17 | 5.20 | cytoplasm |
| PeHDAC3 | Peq001349 | 1149 | 382 | 41765.25 | 5.64 | cytoplasm |
| PeHDAC4 | Peq019669 | 1101 | 366 | 39328.17 | 6.70 | cytoplasm |
| DcaHDAC1 | Dca022379 | 945 | 314 | 34352.98 | 6.49 | cytoplasm |
| DcaHDAC2 | Dca008247 | 2058 | 685 | 76108.86 | 5.06 | cytoplasm |
| DcaHDAC3 | Dca016478 | 1149 | 382 | 41545.88 | 5.44 | cytoplasm |
| DcaHDAC4 | Dca023703 | 1695 | 564 | 62307.71 | 6.31 | nucleus |
| PaHDAC1 | PAXXG039480 | 1338 | 445 | 48369.01 | 6.09 | chloroplast |
| PaHDAC2 | PAXXG172060 | 2034 | 677 | 75165.96 | 5.04 | cytoplasm |
| PaHDAC3 | PAXXG036310 | 1149 | 382 | 41699.15 | 5.57 | cytoplasm |
| PaHDAC4 | PAXXG047590 | 1668 | 555 | 61665.92 | 6.39 | nucleus |
| CeHDAC1 | CETC012070 | 1254 | 417 | 45581.22 | 6.31 | chloroplast |
| CeHDAC2 | CETC014389 | 1251 | 416 | 46375.91 | 5.74 | cytoplasm |
| CeHDAC3 | CETC021136 | 1149 | 382 | 41606.93 | 5.35 | cytoplasm |
| CeHDAC4 | CETC014387 | 1911 | 636 | 70656.73 | 5.36 | cytoplasm |
| CeHDAC5 | CETC008385 | 1698 | 565 | 62709.18 | 6.15 | chloroplast |
| GeHDAC1 | GETC024874 | 2037 | 678 | 75732.99 | 5.09 | cytoplasm |
| GeHDAC2 | GETC008088 | 1686 | 561 | 62576.17 | 8.48 | cytoplasm |
| DchHDAC1 | KAH0460612.1 | 1071 | 356 | 39345.05 | 8.69 | chloroplast |
| DchHDAC2 | KAH0464492.1 | 2058 | 685 | 76286.12 | 5.13 | cytoplasm |
| DchHDAC3 | KAH0470649.1 | 1149 | 382 | 41607.11 | 5.59 | cytoplasm |
| DchHDAC4 | KAH0469025.1 | 1773 | 590 | 65415.18 | 6.21 | cytoplasm |
| DnHDAC1 | KAI0507892.1 | 1335 | 444 | 48050.85 | 6.09 | chloroplast |
| DnHDAC2 | KAI0516625.1 | 2058 | 685 | 76154.90 | 5.05 | cytoplasm |
| DnHDAC3 | KAI0531072.1 | 879 | 292 | 31496.40 | 5.27 | cytoplasm |
| VpHDAC1 | VPTC009343 | 1476 | 491 | 53705.60 | 8.60 | chloroplast |
| VpHDAC2 | VPTC011565 | 2085 | 694 | 77383.33 | 5.03 | cytoplasm |
| VpHDAC3 | VPTC021354 | 1068 | 355 | 42099.79 | 5.84 | cytoplasm |
| VpHDAC4 | VPTC002232 | 1692 | 563 | 62158.41 | 6.37 | nucleus |

To better understand the chromosome distribution of ASDACs on orchid genome, chromosome map of DchHDACs and DnHDACs were constructed based on the annotation information of the *D. chrysotoxum* and *D. nobile* genome (**Figure** 2). The results showed that among 4 DchHDACs, DchHDAC1 was mapped onto the Chr 10, DchHDAC2 was mapped onto the Chr 7, DchHDAC3 was mapped onto the Chr 1, and DchHDAC4 was mapped onto the Chr 3. Moreover, among 3 DnHDACs, DnHDAC1 was localized to the Chr 10, DnHDAC2 was localized to the Chr 7, and DnHDAC3 was localized to the Chr 1.

**FIGURE 1.**
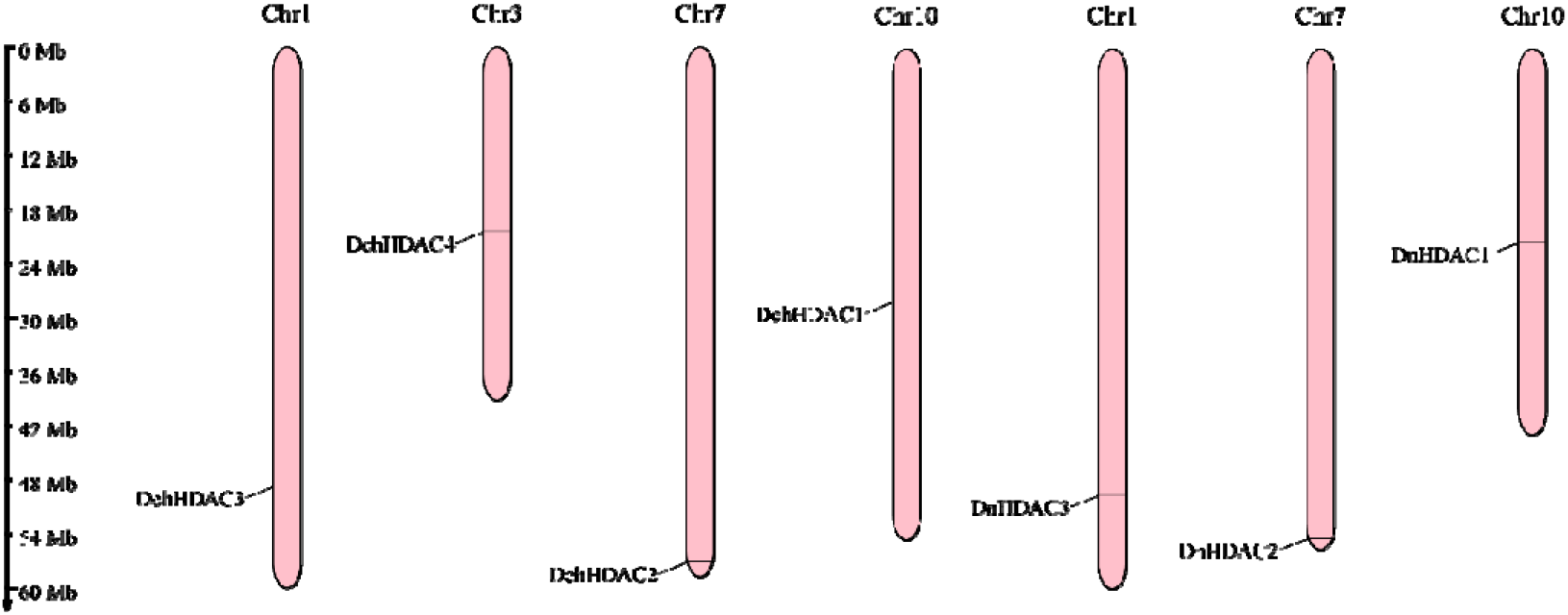
Chromosome distribution of ASDAC proteins of *D. chrysotoxum* and *D. nobile* genome. The chromosome number is shown at the top of each chromosome. Gene position is indicated on the left of the figure.

**FIGURE 2.**
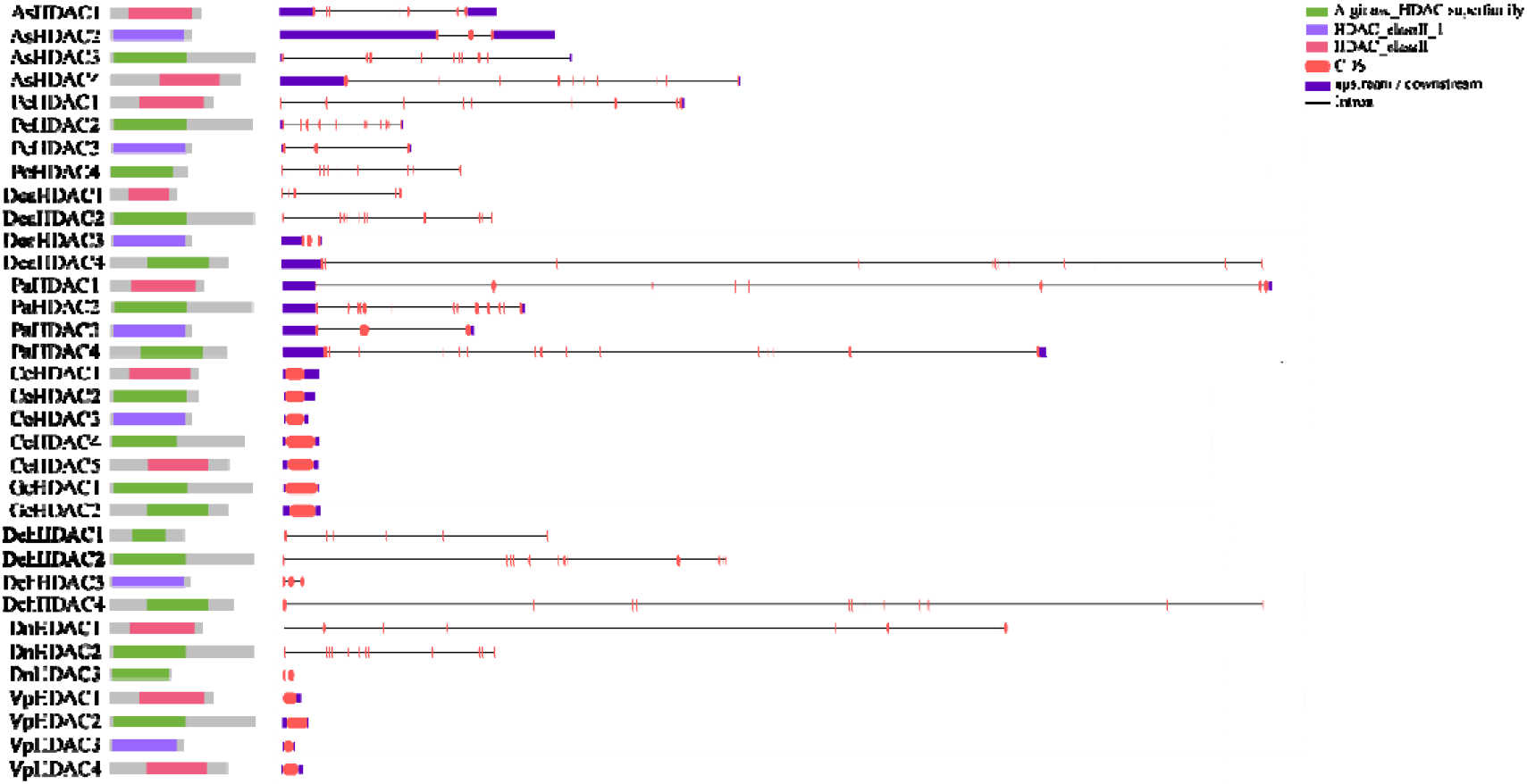

### 3.2 Further Validation of Orchid *ASDACs*

Based on the multiple sequence alignments, we found that *AsHDAC1, PeHDAC1, DcaHDAC1, PaHDAC1, CeHDAC1, DchHDAC1, DnHDAC1*, and *VpHDAC1* are highly homologous with rice and *A. thaliana ASDACs*, and the remaining orchid *ASDACs* have low homology with *OsHDAC10* and *AtHDAC14* (**Supplementary Table S2**). Moreover, in order to exclude incomplete sequence, multiple sequence alignments of the orchid ASDAC domains are performed in **Figure 2A**. The results revealed DchHDAC1 and DnHDAC3 are incomplete in the amino acid sequence at the functional HDAC domain, suggesting the two orchid ASDAC proteins may be inactive. Therefore, we preliminarily speculate that *AsHDAC1, PeHDAC1, DcaHDAC1, PaHDAC1, CeHDAC1, DnHDAC1*, and *VpHDAC1* are homologs of rice and *A. thaliana ASDACs*.

To further understand the evolutionary relationships of orchid *ASDACs*, the phylogenetic tree, motif distribution, and gene structure of orchid *ASDAC* members were analyzed. We constructed a maximum-likelihood (ML) phylogenetic tree using the protein sequences of twenty-five species (**Figure 3A, Supplementary Table S3 and S4**). *ASDAC* homologs were found in various taxa, including archaea, algae, mosses, ferns, and spermatophyte. The phylogenetic tree could be divided into two clades, with strong bootstrap value support. *AsHDAC1, PeHDAC1, DcaHDAC1, PaHDAC1, CeHDAC1, DnHDAC1*, and *VpHDAC1* clustered together with various taxa *ASDACs* (including *OsHDAC10* and *AtHDAC14*), while the other 25 orchid *ASDACs* clustered together separately. Hence, we further speculate that *AsHDAC1, PeHDAC1, DcaHDAC1, PaHDAC1, CeHDAC1, DnHDAC1*, and *VpHDAC1* were potential orchid *ASDACs*, and the remaining 25 members were recognized as *ASDAC*-like genes that belonged to another classes.

**FIGURE 3.**
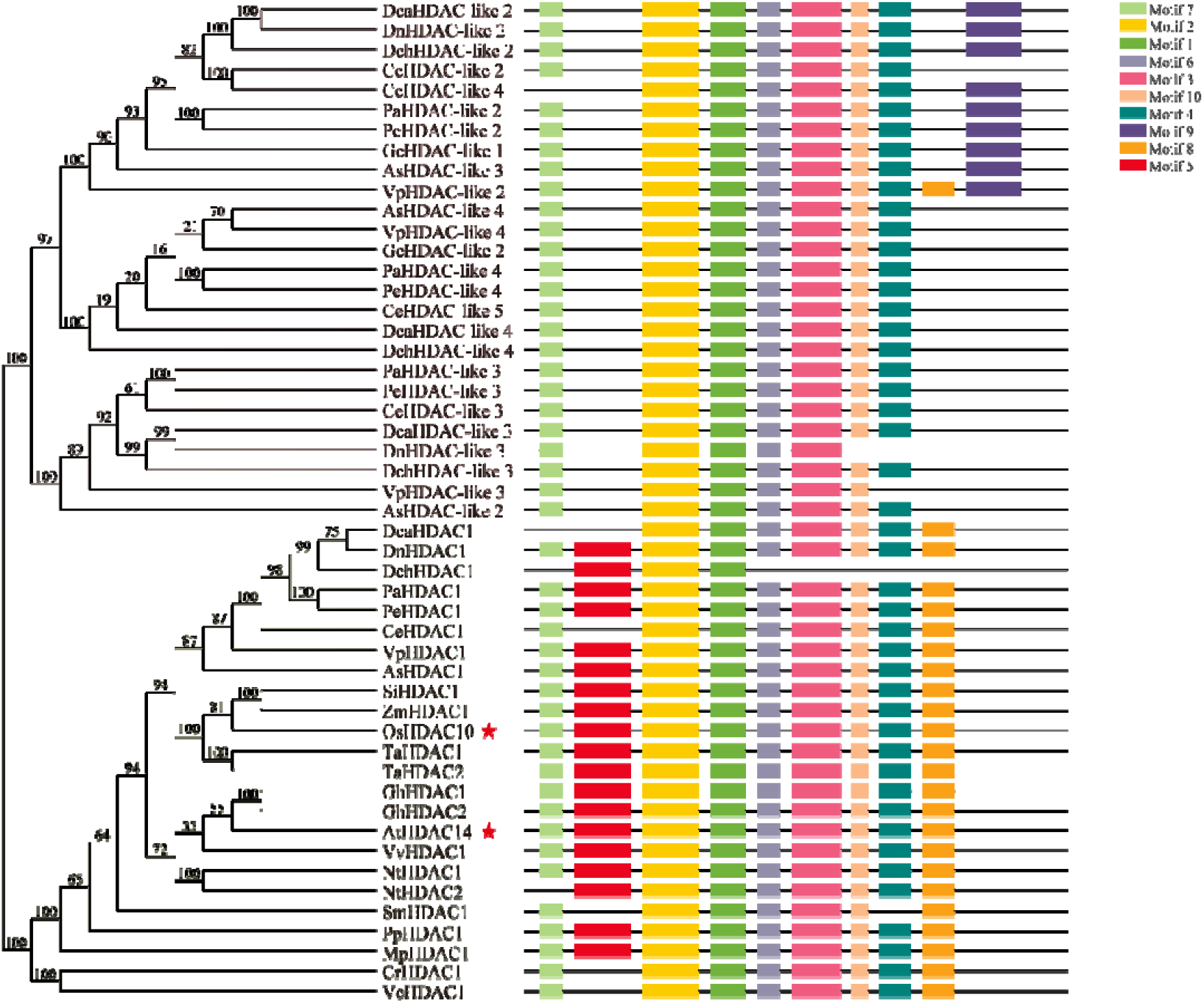
Phylogenetic relationship and motif distribution of orchid *ASDAC* and *ASDAC*-like genes: (A) Phylogenetic tree of the ASDAC family proteins from various taxa. The red pentacles represent the functionally confirmed *ASDACs*. (B) The conserved motif patterns for each gene. Different motifs are highlighted with various colored boxes. *MtHDACs* (*Methanosaeta thermophila*), *MhHDACs* (*Methanothrix harundinacea*), *MsHDACs* (*Methanothrix soehngenii*), *VcHDACs* (*Volvox carteri*), *CrHDACs* (*Chlamydomonas reinhardtii*), *MpHDACs* (*Marchantia polymorpha*), *PpHDACs* (*Physcomitrella patens*), *SmHDACs* (*Selaginella moellendorffii*), *SiHDACs* (*Setaria italica*), *ZmHDACs* (*Zea mays*), *TaHDACs* (*Triticum aestivum*), *VvHDACs* (*Vitis vinifera*), *GhHDACs* (*Gossypium hirsutum*), *NtHDACs* (*Nicotiana tabacum*).

From gene structure results, except for the number of introns was 0 in *ASDAC* and *ASDAC*-like genes of *C. ensifolium, G. elata*, and *V. planifolia*, the quantity of introns in *ASDAC* and *ASDAC*-like genes of *A. shenzhenica, P. equestris, D. catenatum, P. aphrodite, D. chrysotoxum*, and *D. nobile* ranged from 2 to 16 (**Figure 3C**). Combined exon-intron structural and phylogenetic analysis, most structurally similar orchid *ASDAC* or *ASDAC*-like genes clustered together in the same clades of phylogenetic trees. For example, *AsHDAC1, PeHDAC1, DcaHDAC1, PaHDAC1*, and *DnHDAC1* of the same clades were found to be comprised of 4-8 introns. Motif is a short sequence of relatively conserved features shared by genes^[31]^. In motif analysis results, a total of 10 distinct motifs, named motif 1 to motif 10, were detected in *ASDAC* genes from various taxa (**Figure 3B and 4**). The lengths of these conserved motifs varied from 22 to 50 amino acids (**Figure 4**). All various taxa *ASDACs* contained motif 9 besides archaea *ASDACs*, indicating the motif might have specific roles. Moreover, the same cluster’s orchid *ASDAC* or *ASDAC*-like proteins generally contained similar motifs. These results of gene structure and motif analysis verified the closeness of the phylogenetic tree in orchid, suggesting that *AsHDAC1, PeHDAC1, DcaHDAC1, PaHDAC1, CeHDAC1, DnHDAC1*, and *VpHDAC1* within the same clade may play similar functional roles in melatonin biosynthesis.

**FIGURE 4.**
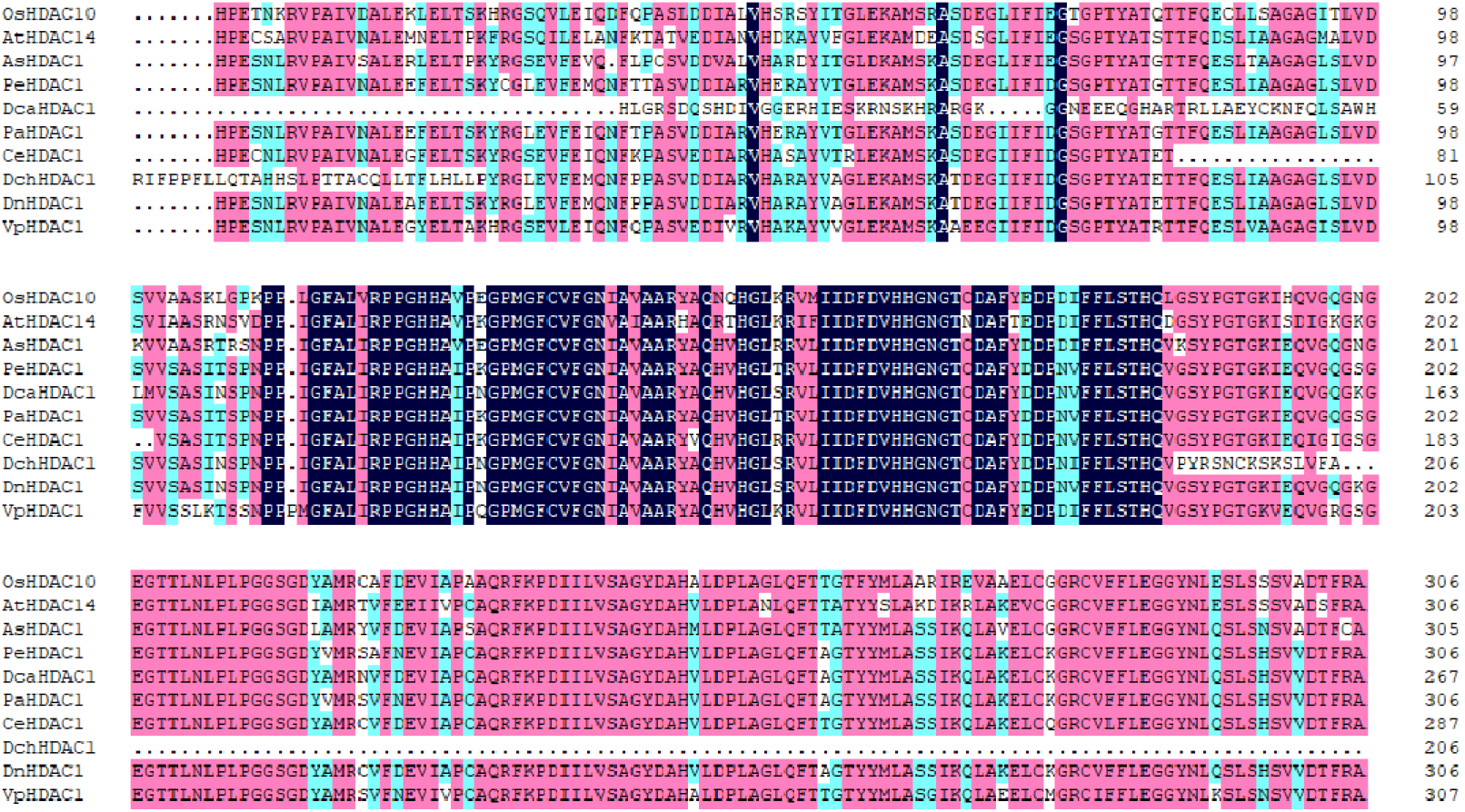

Commonly, the Ka/Ks ratio can show Ka/Ks >1 (positive selection), Ka/Ks <1 (negative or purifying selection), and Ka/Ks =1 (neutral selection)^[32]^. In this study, based on the phylogenetic relationship, the Ka/Ks values of ASDAC proteins from lower plant to higher plant were calculated to evaluate the driving force underlying the orchid *ASDAC* gene’s evolution. The results showed that Ka/Ks values of gene pairs ranged from 0.0031 ∼ 0.7798, suggesting that orchid *ASDACs* were under purifying selection during the evolution process (**Figure 5, Supplementary Table S5**).

**FIGURE 5.**
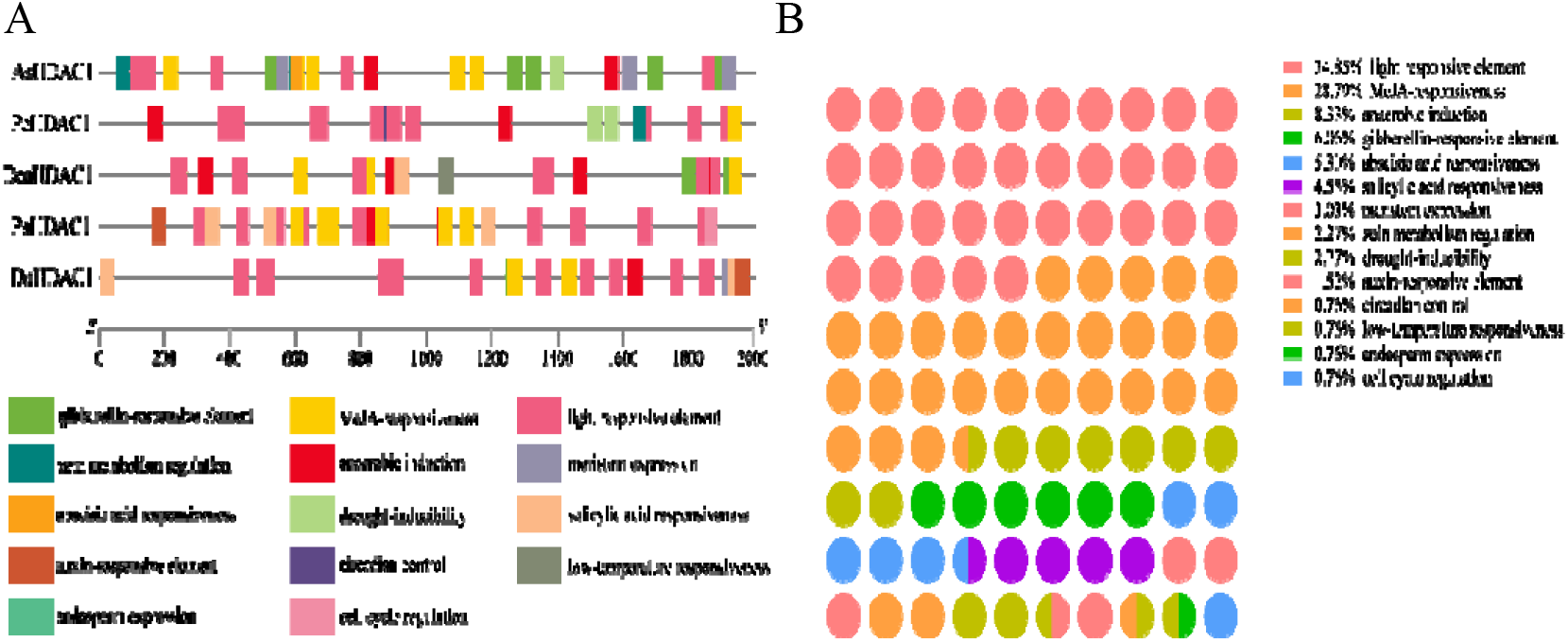
*Cis*-Active regulatory elements in the promoters of orchid *ASDAC* genes (A) and their proportions (B). The different colored blocks represent the diverse types of *cis*-elements and their locations in each gene.

### 3.3 Functional Analysis of Orchid *ASDAC* Genes

To better investigate the potential function of orchid *ASDACs*, we analyzed their *cis*-active regulatory elements and expression patterns (**Figure 6A**). However, *CeHDAC1* and *VpHDAC1* could not detected the *cis*-elements on account of a short sequence upstream of start codon (ATG). The promoter regions of *AsHDAC1, PeHDAC1, DcaHDAC1, PaHDAC1*, and *DnHDAC1* detected 14 *cis*-elements, containing plant growth and development responsive elements, phytohormone responsive elements, and stress responsive elements.

**FIGURE 6.**
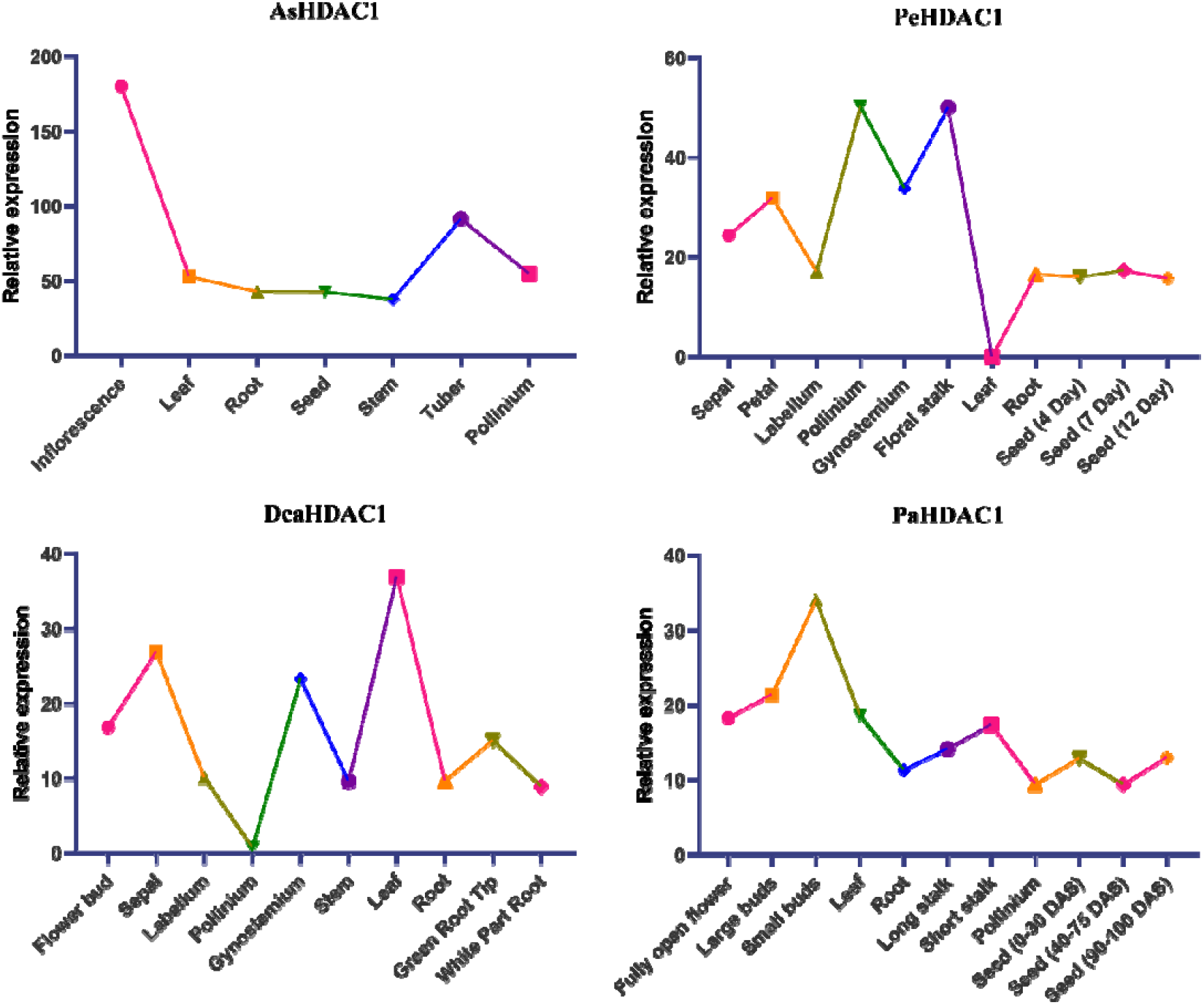
Expression patterns of *A. shenzhenica, P. equestris, D. catenatum*, and *P. aphrodite ASDAC* gene in different tissues.

Most phytohormones can regulate plant growth and development. Among the 14 *cis*-regulatory elements, phytohormone-related elements (including gibberellin, methyl jasmonate, abscisic acid, auxin, and salicylic acid) were the largest category (46.22 %) (**Figure 6B**). The plant growth and development responsive elements were also found in the promoter regions of orchid *ASDAC* genes (**Figure 6B**). Furthermore, most orchid *ASDACs* were expressed in the roots, stems, leaves, flowers, and seeds (**Figure 7, Supplementary Table S6**). Among them, *AsHDAC1, PeHDAC1*, and *PaHDAC1* exhibited a relatively higher transcript accumulation in the flowers. Similarly, *VpHDAC1* and *DcaHDAC1* was expressed at relatively high levels in the leaves. However, *PeHDAC1* was not expressed in the leaves. The expression of *AsHDAC1, PeHDAC1*, and *PaHDAC1* were relatively low in the roots and seeds. The expression of *AsHDAC1, DcaHDAC1*, and *VpHDAC1* were also relatively low in the stems. *CeHDAC1* were only expressed in pooled 4 stage of bud and mature flower based on public databases. These results indicated that orchid *ASDAC* genes could response to various hormones to regulate the growth and development of orchids.

Among the 5 orchid ASDACs, the stress-related responsive elements mainly consisted of light responsive element, anaerobic induction, drought-inducibility, and low-temperature responsive element, accounted for the second largest proportion (46.21 %) (**Figure 6B**). Moreover, all genes contained the light responsive element, accounted for 34.85 % of the total. *AsHDAC1, DcaHDAC1, PaHDAC1*, and *VpHDAC1* were expressed in the leaves (**Figure 7**). These results demonstrating that orchid *ASDAC* genes might play important roles in plant photoperiod regulation. And virtually all promoter of genes contained more than two stress-related elements, suggesting that the elements could be involved in orchid *ASDACs* expression regulation in response to multiple abiotic stresses.

## 4. DISCUSSION

Orchidaceae is a conspicuous family with highly ornamental and medical value^[3]^. Additionally, orchids are distributed in almost every habitat on the Earth and face various environmental stresses. As a novel phytohormone, melatonin not only regulate plant growth and development, but also respond to diverse environmental stresses. The effects of melatonin on orchids have remained poorly understood. ASDAC is the only reverse enzyme in melatonin biosynthesis and plays an important role in regulating the balance of melatonin in plants (**Figure 8**)^[14]^. However, *ASDAC* genes have been merely identified in rice and *A. thaliana*^[14]^. In orchids, the role of *ASDACs* in regulating melatonin remains unclear.

**FIGURE 8.**
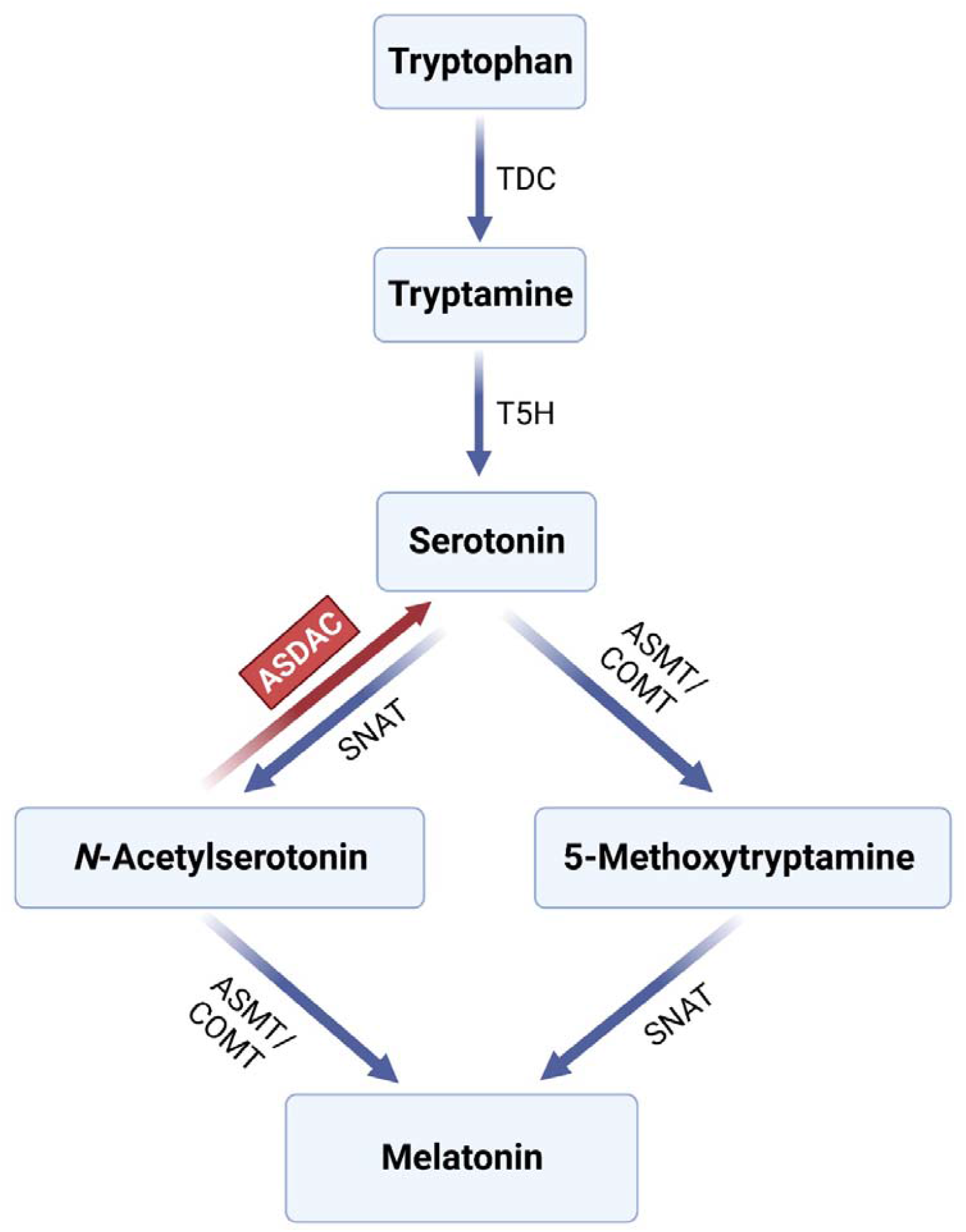
The biosynthesis of melatonin in plants. ASDAC (N-acetylserotonin deacetylase) is an only reverse enzyme of melatonin biosynthesis pathway; TDC, tryptophan decarboxylase; T5H, tryptamine 5-hydroxylase; SNAT, serotonin N-acetyltransferase; ASMT, acetylserotonin methyltransferase; COMT, caffeic-O-methyltransferase.

For our study, a total of 7 candidate *ASDAC* genes were recognized from nine species of Orchidaceae. The 7 orchid *ASDACs* were highly homologous to rice and *A. thaliana ASDACs*. Furthermore, *AsHDAC1, PeHDAC1, DcaHDAC1, PaHDAC1, CeHDAC1, DnHDAC1*, and *VpHDAC1* clustered together with *OsHDAC10* and *AtHDAC14*. These findings specifically support a role for orchid *ASDACs* in melatonin biosynthesis, deacetylates N-acetylserotonin to serotonin.

The analysis of gene structure, motif distribution, and phylogenetic tree can provide important clue for the evolutionary relationship of genes. Most of the orchid *ASDAC* or *ASDAC*-like genes with similar introns and motifs cluster in the same branch of the phylogenetic tree, suggesting the distribution pattern of exons/introns and motifs were strongly related to phylogeny on an evolutionary basis. Moreover, apart from the identification of orchid *ASDACs*, we also recognized family members of *ASDAC* in archaea, algae, mosses, ferns, and spermatophyta with one to three members in each species. These results indicated the highly evolutional conservation of ASDAC protein functions from lower to higher plants.

The ASDAC and serotonin N-acetyltransferase (SNAT) both regulate the conversion between serotonin and N-acetylserotonin in melatonin biosynthesis pathway^[33]^. ASDAC catalyze N-acetylserotonin into serotonin, whereas SNAT catalyze serotonin into N-acetylserotonin. An interesting feature of ASDAC and SNAT is their subcellular localization, both *OsHDAC10, AtHDAC14*, and plant *SNATs* shows chloroplasts localization^[14, 34]^. In this study, the prediction of protein subcellular location indicates *PaHDAC1, CeHDAC1, DnHDAC1*, and *VpHDAC1* located in chloroplasts as well. Overall, it suggests a strict regulation for melatonin synthesis in specific organelles or tissues.

Additionally, it is worth noting that ASDAC families had fewer members in *G. elata* than in the other eight species. Furthermore, all *ASDACs* of *G. elata* have low homology and not cluster together with rice and *A.thaliana ASDACs. G. elata* has a fully mycoheterotrophic lifestyle that is different from *A. shenzhenica, P. equestris, D. catenatum, P. aphrodite, C. ensifolium, D. chrysotoxum, D. nobile*, and *V. planifolia*^[35]^. It has been reported that *G. elata* displayed the largest extent of gene family contraction and undergone extensive gene loss events^[35, 36]^. Therefore, it is speculated that *ASDAC* gene family was lost in *G. elata*. Recently, the chromosome-scale assembled genomes of *Platanthera guangdongensis* was reported, a fully mycoheterotrophic orchid^[37]^. We also retrieve the putative *ASDAC* genes from *P. guangdongensis* (**Supplementary Table S7**). Similarly, the *ASDACs* of *P. guangdongensis* share low sequence identity with *OsHDAC10* and *AtHDAC14*. These results further supported that *ASDAC* genes were lost in the holomycoheterotrophic orchids.

Different *cis*-acting elements were observed in the promoter regions of orchid *ASDAC* genes, and provide an understanding of its role in various biological processes. The upstream region of all orchid *ASDACs* contain plentiful of light-responsive elements. Moreover, some orchid *ASDACs* possess the *cis*-acting elements related to abiotic stress, such as drought and low-temperature responsive elements. Leaves can exhibit biotic stress responses. Our results showed that orchid *ASDACs* were expressed in the leaves. Combined with the function of melatonin in abiotic stresses including light, drought and low temperature stress, orchid *ASDACs* may play an important role in melatonin response to various stress^[13, 38]^.

In plants, phytohormone such as gibberellins, jasmonate, ethylene, cytokinins, abscisic acid, auxin, and salicylic acid play important roles in regulating growth and development^[39, 40]^. In this study, the auxin, gibberellins, abscisic acid, methyl jasmonate, and salicylic acid responsive elements were found in the promoter regions of most orchid *ASDACs*. Furthermore, orchid *ASDACs* were widely expressed in roots, stems, leaves, flowers, and seeds. Combined with the *cis*-acting element and tissue expression analysis, orchid *ASDAC* genes could regulate the growth and development of orchids in response to various hormones. Additionally, melatonin regulated plant growth and development, as well as mediated the interaction with plant hormone^[41]^. The results further indicates that orchid *ASDAC* genes are involved in the cross-talk between melatonin and other plant hormone in regulating orchids growth and development.

orchids are important plants due to their great value in ornamental. The high expression of orchid *ASDACs* in the flowers indicated that *ASDAC* genes could be pivotal in flower growth and development. Previous researches have shown that melatonin regulates flowering in a dose-dependent manner. The increased melatonin levels at a suitable range resulted in more flowering^[42]^. When the concentration of melatonin was too high, the flowering rate was decreased and the flowering time was delayed^[42]^. The data suggested that *ASDAC* genes were expressed at a high level to maintain melatonin balance. In this way, melatonin may be maintained at an optimal level, which is essential for flower growth and development.

## 5. CONCLUSION

In summary, genome-wide identification and analyses of the *ASDAC* gene family were firstly conducted in Orchidaceae. A total of 7 *ASDAC* genes were identified in orchids. Phylogenetic analysis revealed that the 7 orchid *ASDACs* clustered together with the functionally confirmed *OsHDAC10* and *AtHDAC14*, as well as the exon/intron distributions and conserved motif compositions were strongly related to phylogeny. Furthermore, based on the phylogenetic relationship, the Ka/Ks ratios of *ASDAC* gene pairs from lower plant to higher plant were less than one, suggested that orchid *ASDAC* genes have undergone purifying selection during the evolution process. Analysis results of *cis*-acting elements showed that the promoter regions of orchid *ASDAC* genes contained plant growth and development, phytohormone, and stress responsive elements. Moreover, most orchid *ASDACs* were expressed in the roots, stems, leaves, flowers, and seeds. Combined *cis*-acting element and tissue expression analysis, suggesting orchid *ASDAC* genes are involved in melatonin regulation of the growth and development, as well as melatonin responds to various stresses in orchids. In short, this study provides a reference for further exploring the function of orchid *ASDAC* gene in melatonin biosynthesis and the role of melatonin in orchids.

